# *miR-9a* regulates levels of both *rhomboid* mRNA and protein in the early *Drosophila melanogaster* embryo

**DOI:** 10.1101/2021.07.12.452096

**Authors:** Lorenzo Gallicchio, Sam Griffiths-Jones, Matthew Ronshaugen

## Abstract

MicroRNAs have subtle and combinatorial effects on the expression levels of their targets. Studying the consequences of a single microRNA knockout often proves difficult as many such knockouts exhibit phenotypes only under stress conditions. This has led to the hypothesis that microRNAs frequently act as buffers of noise in gene expression. Observing and understanding buffering effects requires quantitative analysis of microRNA and target expression in single cells. To this end, we have employed single molecule fluorescence *in situ* hybridization, immunofluorescence, and high-resolution confocal microscopy to investigate the effects of *miR-9a* loss on the expression of the serine-protease rhomboid in *Drosophila melanogaster* early embryos. Our single-cell quantitative approach shows that *rhomboid* mRNA exhibits the same spatial expression pattern in WT and *miR-9a* knockout embryos, although the number of mRNA molecules per cell is higher when *miR-9a* is absent. However, the level of rhomboid protein shows a much more dramatic increase in the *miR-9a* knockout. Specifically, we see accumulation of rhomboid protein in *miR-9a* mutants by stage 5, much earlier than in WT. The data therefore show that *miR-9a* functions in the regulation of *rhomboid* activity by both inducing mRNA degradation and inhibiting translation in the blastoderm embryo. Temporal regulation of neural proliferation and differentiation in vertebrates by *miR-9* is well-established. We suggest that *miR-9* family microRNAs are conserved regulators of timing in neurogenic processes. This work shows the power of single-cell quantification as an experimental tool to study phenotypic consequences of microRNA mis-regulation.

## Introduction

The study of development in *Drosophila melanogaster* embryos, larvae, and adults has provided an extremely important model for the study of microRNA (miRNA) biogenesis and function (Matranga *et al*. 2005; Rand *et al*. 2005; Okamura *et al*. 2007). MicroRNAs are short ∼22 nucleotide long, single-stranded, endogenous RNAs found in animals and plants (Bartel 2004; Kozomara *et al*. 2019). MicroRNAs regulate gene expression post-transcriptionally by recruiting the RNA-induced silencing complex (RISC) and then binding to specific sequences on target mRNA molecules, usually in their 3’UTR. The binding of the miRNA-RISC triggers repression of translation, deadenylation, and/or degradation of the target mRNA (Valencia-Sanchez *et al*. 2006). It is estimated that the majority of animal mRNAs are targeted by miRNAs (Friedman *et al*. 2009; Agarwal *et al*. 2015). An intriguing debate has arisen regarding the phenotypic consequences of miRNA mis-regulation, with GOF (gain of function) and LOF (loss of function) studies in different organisms finding that they act as either minor modulators or key regulators of gene expression (Miska *et al*. 2007; Alvarez-Saavedra and Horvitz 2010; Chen *et al*. 2014).

In many cases, individual effects of miRNAs on the expression of a target are relatively small (Miska *et al*. 2007; Alvarez-Saavedra and Horvitz 2010; Chen *et al*. 2019a). In addition, each miRNA may target hundreds of different transcripts, and many different miRNAs have been found to act on the same targets (Peter 2010). It is therefore expected that a high degree of quantitative precision is required to determine specific effects of miRNAs on gene expression. Indeed, a complete understanding of miRNA function will only come from a precise quantitative analysis of miRNA activity at the single cell level. Single cell studies of miRNA effects on gene regulation may provide insight into mis-regulation phenotypes that are not apparent at a tissue or organism level (Miska *et al*. 2007; Alvarez-Saavedra and Horvitz 2010). It has also been observed that the phenotypic effects of miRNA mutation or mis-regulation are sometimes only revealed under particular conditions (e.g. dietary or temperature stresses) (Li *et al*. 2009; Kennell *et al*. 2012). For example, flies lacking *miR-14* are more sensible to salt stress compared to WT, while flies lacking *miR-7* present abnormal expression of the proteins Yan and Ato only under temperature fluctuations (Xu *et al*. 2003; Li *et al*. 2009). Such stress-dependent miRNA phenotypes have also been observed in other organisms such as mouse and zebrafish (Van Rooij *et al*. 2007; Flynt *et al*. 2009). Thus, the phenotypic consequences of miRNA mis-regulation may be subtle and cryptic.

The *mir-9* miRNA family is highly conserved in bilaterians and is a good example of a miRNA that can exhibit both subtle and strong phenotypes (Coolen *et al*. 2013). Experiments in a variety of vertebrate models show conservation of *mir-9* expression and function in neurogenesis and neuronal progenitor proliferation. Over-expression of *mir-9* in zebrafish embryos (Leucht *et al*. 2008), mouse embryonic cortex (Zhao *et al*. 2009) and chicken spinal cord (Otaegi *et al*. 2011) leads to a reduction of the number of proliferating progenitors, similarly to the observed effects in *Drosophila* (Li *et al*. 2006). Also common to these studies is the observation that *mir-9* alteration (both loss and gain of function) results in a quite mild phenotype (Shibata *et al*. 2011). This supports the idea that *mir-9* is not a biological switch that allows the cell to adopt a certain fate, but a control factor to maintain a proper development trajectory, possibly acting as a key component of a feedback control system. *mir-9* dysfunction has been associated with a number of human pathologies, including various kinds of cancer and neurodegenerative disorders (Coolen *et al*. 2013; He *et al*. 2017; Chen *et al*. 2019b; Khafaei *et al*. 2019). In medulloblastomas (a paediatric brain cancer) tumour cells appear to have a decreased expression of *mir-9*, while in a subclass of glioblastoma (an aggressive adult brain cancer) tumour cells express *mir-9* at a higher level (Ferretti *et al*. 2009; Kim *et al*. 2011). *mir-9* has been found to have a role also in cancers not directly related with the nervous system, in which it may act as an oncogene or a tumour suppressor (Coolen *et al*. 2013). These dual roles and opposite effects, combined with observations of subtle and cryptic phenotypes, has led to a model where miRNAs act to control or modulate the dynamics of biological processes, and not as biological switches themselves.

Many studies have focused on *miR-9a* as a modulator of the specification and number of *Drosophila* sensory organ precursor (SOP) cells, a key neuronal cell type that emerges during embryonic stage 10 (Li *et al*. 2006; Cassidy *et al*. 2013). At embryonic stage 5, *miR-9a* is expressed in the dorsal ectoderm and in the neuroectoderm: the germ layer in which the future neuronal precursor cells will form (Fu *et al*. 2014; Gallicchio *et al*. 2021). It is possible that *miR-9a* functions as a modulator of the genes required for proper ectoderm and neuroectoderm specification. This early expression is reminiscent of *miR-1*, a miRNA involved in mesoderm specification and muscle development, which is also expressed during early embryogenesis exclusively in the presumptive mesoderm (Sokol and Ambros 2005). Moreover, it has been suggested that both miRNAs might respond to the dorsal TF gradient that activates and inhibits expression of genes involved in establishing germ layers (Biemar *et al*. 2006). It is reported that *miR-9a* KO flies show defects on the wing margin (Li *et al*. 2006) and an homozygous KO for *miR-1* causes lethality in second instar larvae, which die immobilized and with abnormal musculature (Sokol and Ambros 2005). Nevertheless, no differences in germ layer specification during embryogenesis have ever been observed in either *miR-9a* or *miR-1* mutant (Fu *et al*. 2014). This is perhaps not surprising, as multiple miRNAs often function redundantly, and it is rare that a specific biological process is strongly affected when a single miRNA is knocked out (Liufu *et al*. 2017). However, when *miR-1* and *miR-9a* are mutated together dramatic effects on embryonic development are observed (Fu *et al*. 2014). The double knockout displayed an ectopic overexpression of *rhomboid* (*rho*), a dorsal target gene expressed in the neuroectoderm, and a failure of gastrulation (Fu *et al*. 2014). *rho* possesses two *miR-9a* binding sites on its 3’UTR, indicating that *miR-9a* might directly regulate *rho* mRNA degradation and/or translation.

We were therefore motivated to study *rho* expression in single cells and compare quantification of mRNA number and protein levels between WT and *miR*-*9a* KO embryos in the establishment of the embryonic domain of *rho* mRNA and protein. Using high resolution confocal microscopy coupled with multiplex smFISH and IF we examined expression domains, transcription dynamics and protein accumulation at the single cell level in whole mount developing *D. melanogaster* embryos. In *miR-9a* KO mutants, we observed an increase in both *rho* mRNA number per cell and Rho protein expression. We therefore conclude that *rho* is directly targeted by *miR-9a*. Together, these results show that single-cell analysis and quantification is a powerful approach to study miRNA function on target gene expression.

## Materials and Methods

### Fly stocks, embryo collection, and fixing and larval dissection

Flies were grown at 25 or 18°C. Embryos were collected after ∼20 h and fixed in 1 V heptane + 1 V 4% formaldehyde for 30 min shaking at 220 rpm. The embryos were then washed and shaken vigorously for one minute in 100% methanol. Fixed embryos were stored in methanol at −20°C. Larvae were dissected in 1× PBS, carcasses were fixed in 1 V 1× PBS + 1 V 10% formaldehyde for ∼1 h, washed with methanol, and stored in methanol at −20°C. Genotypes used for this study are: W [1118], (from Bloomington Drosophila Resource Centre) and *miR-9a*^E39^ mutants (Li *et al*. 2006) generously gifted by the Fen-Biao Gao lab.

### Probe design, smFISH, and Immunofluorescence

We applied an inexpensive version (Tsanov *et al*. 2016; Morales-Polanco *et al*. 2021) of the conventional smFISH protocol in *Drosophila* (Trcek *et al*. 2017). Primary probes were designed against the mature *rho* mRNA (*rhomboid_e*), the first *rho* intron (*rhomboid_i*) and a genomic region flanking the *mir-9a* gene locus using the Biosearch Technologies Stellaris probe Designer (version 4.2). All sequences were obtained from FlyBase. To the 5′ end of each probe was added the Flap sequence CCTCCTAAGTTTCGAGCTGGACTCAGTG. Multiple secondary probes that are complementary to the Flap sequence were tagged with fluorophores (CAL Fluor Orange 560, CAL Fluor Red 610, Quasar 670) to allow multiplexing. Probes sequences are reported in Supplementary Data. For Immunofluorescence we used the following antibodies: mouse anti-Dorsal (Developmental Studies Hybridoma Bank #AB_528204) at 1:100, mouse anti-Spectrin (Developmental Studies Hybridoma Bank #AB_520473) at 1:100, guinea-pig anti-Rho gently gifted from the Hayashi lab at 1:400 (Ogura *et al*. 2018), goat anti-guinea pig IgG (H + L) Highly Cross-Adsorbed Secondary Antibody Alexa Fluor 555 (Invitrogen #A21435) at 1:500, and goat anti-mouse IgG (H + L) Highly Cross-Adsorbed Secondary Antibody Alexa Fluor 488 (Invitrogen #A32723) at 1:500. Further details on reagents used are provided in the Reagents Table.

### Imaging and quantification

Imaging was performed using a Leica SP8 Inverted Tandem Head confocal microscope with LAS X v.3.5.1.18803 software (University of Manchester Bioimaging facility), using 40×, and 100× magnifications. Deconvolution was performed using Huygens Pro v16.05 software. Membrane segmentation was performed on Imaris (version 9.5.0), mRNA molecules and Transcription sites were counted after membrane segmentation on Imaris 9.5.0 using the Cell module. Protein fluorescence levels were measured using FIJI for Macintosh. From each picture, five measurements of background mean intensity were taken. Each single measurement was then adjusted using the formula: integrated density – (area × background mean).

### Data availability statement

Strains and plasmids are available upon request. The authors affirm that all data necessary for confirming the conclusions of the article are present within the article, figures, and tables.

## Results

### *rho* and *mir-9a* are co-expressed in the neurogenic ectoderm

After the identification of Rhomboid (Rho) as an intramembrane serine protease in *Drosophila*, Rho-like proteins have been identified in nearly every metazoan, suggesting a conserved role for the family (Urban *et al*. 2001; Freeman 2014). Although the molecular and cellular function of Rho-like proteins is well established, how their expression is post-transcriptionally regulated has not been examined in detail. We therefore decided to investigate if *miR-9a* and/or *miR-1* could directly regulate *rho* mRNA degradation and/or translation. As *miR-1* is exclusively expressed in the mesoderm (Sokol and Ambros 2005; Fu *et al*. 2014) and *miR-9a* in the dorsal and neurogenic ectoderm (Fu *et al*. 2014; Gallicchio *et al*. 2021) largely overlapping *rho* (Ip *et al*. 1992a), we hypothesize that *miR-9a* might directly target *rho*. We used TargetScan (Agarwal *et al*. 2018) and SeedVicious (Marco 2018) to computationally verify the presence of two potential *miR-9a* binding sites in the *D. melanogaster rho* 3’UTR. *rho* has 2 alternatively polyadenylated transcripts (based on the most recent gene annotation in FlyBase), and the predicted *miR-9a* binding sites are both located in the common 3’UTR region. In addition, we used SeedVicious to verify the presence of *miR-9a* binding sites on Rho orthologs in beetle (*Tribolium castaneum*), worm (*Caenorhabditis elegans*), zebrafish (*Danio rerio*), mouse (*Mus musculus*) and human, and the non-model organisms mosquito (*Anopheles gambie*), butterfly (*Heliconius melpomene*) and mite (*Tetranychus urticae*) (Table 1).

**Table 1.**
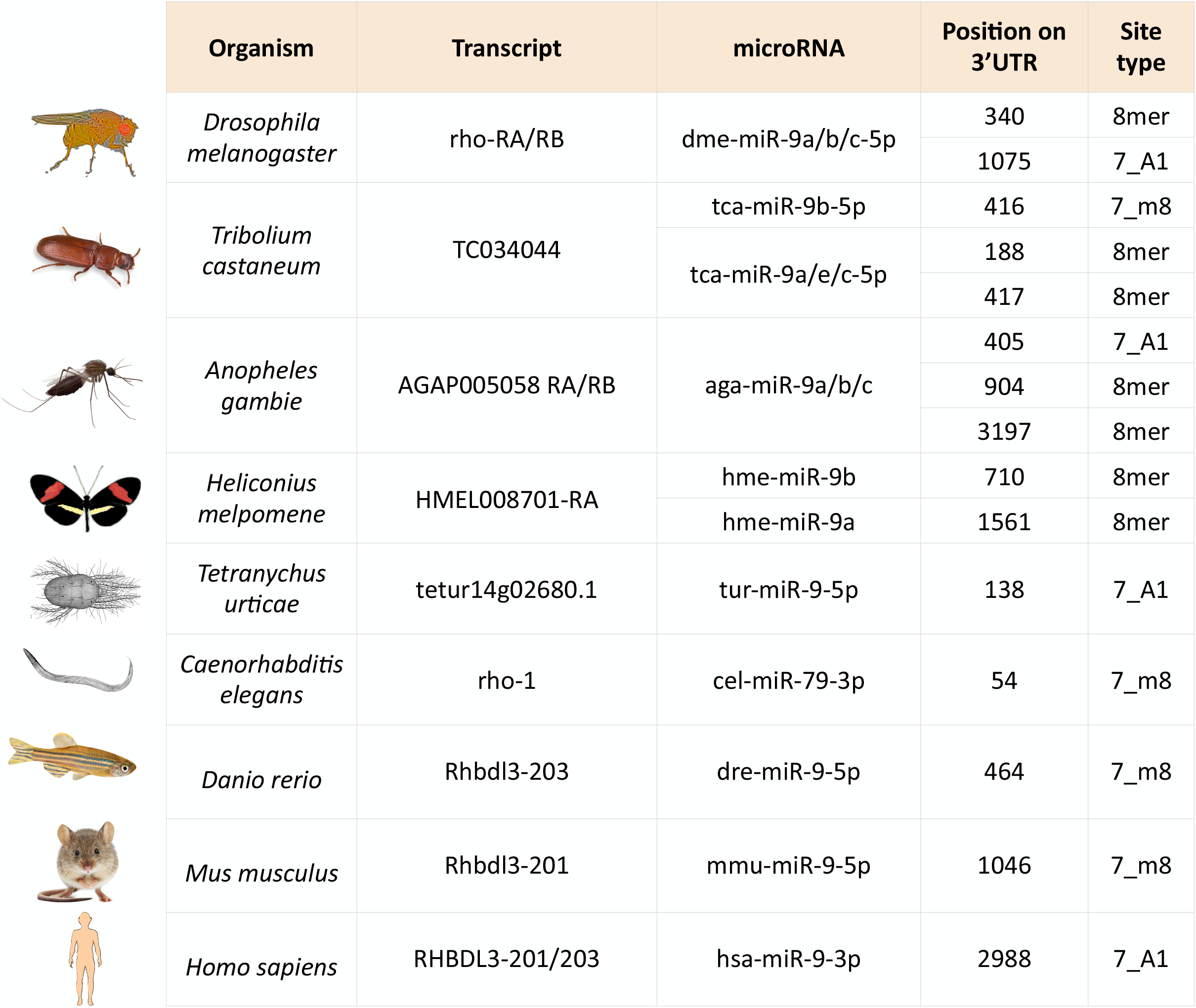

|  | Organism | Transcript | microRNA | Position on 3'UTR | Site type |
| --- | --- | --- | --- | --- | --- |
|  | *Drosophila melanogaster* | rho-RA/RB | dme-miR-9a/b/c-5p | 340 | 8mer |
|  |  |  |  | 1075 | 7_A1 |
|  | *Tribolium castaneum* | TC034044 | tca-miR-9b-5p | 416 | 7_m8 |
|  |  |  | tca-miR-9a/e/c-5p | 188 | 8mer |
|  |  |  |  | 417 | 8mer |
|  | *Anopheles gambiae* | AGAP005058 RA/RB | aga-miR-9a/b/c | 405 | 7_A1 |
|  |  |  |  | 904 | 8mer |
|  |  |  |  | 3197 | 8mer |
|  | *Heliconius melpomene* | HMEL008701-RA | hme-miR-9b | 710 | 8mer |
|  |  |  | hme-miR-9a | 1561 | 8mer |
|  | *Tetranychus urticae* | tetur14g02680.1 | tur-miR-9-5p | 138 | 7_A1 |
|  | *Caenorhabditis elegans* | rho-1 | cel-miR-79-3p | 54 | 7_m8 |
|  | *Danio rerio* | Rhbdl3-203 | dre-miR-9-5p | 464 | 7_m8 |
|  | *Mus musculus* | Rhbdl3-201 | mmu-miR-9-5p | 1046 | 7_m8 |
|  | *Homo sapiens* | RHBDL3-201/203 | hsa-miR-9-3p | 2988 | 7_A1 |

We also employed nascent transcript smFISH to precisely establish the overlap in expression domains of *rho* and the primary transcript of *miR-9a* (pri-*mir-9a*). To identify cells that are actively transcribing *rho*, we designed probes against the first intron of *rho* to detect active transcription sites (TS). As mature miRNAs are too short to be detected via smFISH, we designed probes against ∼1kb of sequence flanking the *mir-9a* hairpin to detect the larger primary transcript. Using multiplex smiFISH, we were able to identify cells that are transcribing both *rho* and *mir-9a* at the same time (Figure 1). As expected, *rho* expressing cells are contained entirely within the *mir-9a* expression domain (Figure 1 A-B). Since it has been widely observed that gene expression patterns are highly dynamic during stage 5 (Reeves *et al*. 2012), we measured membrane introgression to distinguish between stage 5 sub-stages. We find that both *rho* and *mir-9a* expression pattern become more defined at the ventral edge of their expression domain as stage 5 proceeds (Figure 1 C-F). Interestingly, while *rho* expressing cells are generally also expressing *mir-9a*, there are many cells at the ventral edge that are expressing only *mir-9a* (Figure 1 C). As stage 5 progresses the two genes become co-expressed in the same cells, which mark a clear boundary between neurogenic ectoderm and presumptive mesoderm (Figure 1 D). It is therefore possible that the two genes respond differently to the Dorsal gradient, specifically to the repressor *snail*, which has been shown to repress both *mir-9a* and *rho* in the mesoderm (Hemavathy *et al*. 2004; Fu *et al*. 2014). Taken together, the co-expression of *rho* and *mir-9a* and presence of conserved *miR-9a* target sites suggest that *miR-9a* is a strong candidate to target *rho* mRNA during embryogenesis, and that this role may be evolutionally conserved.

**Figure 1.**
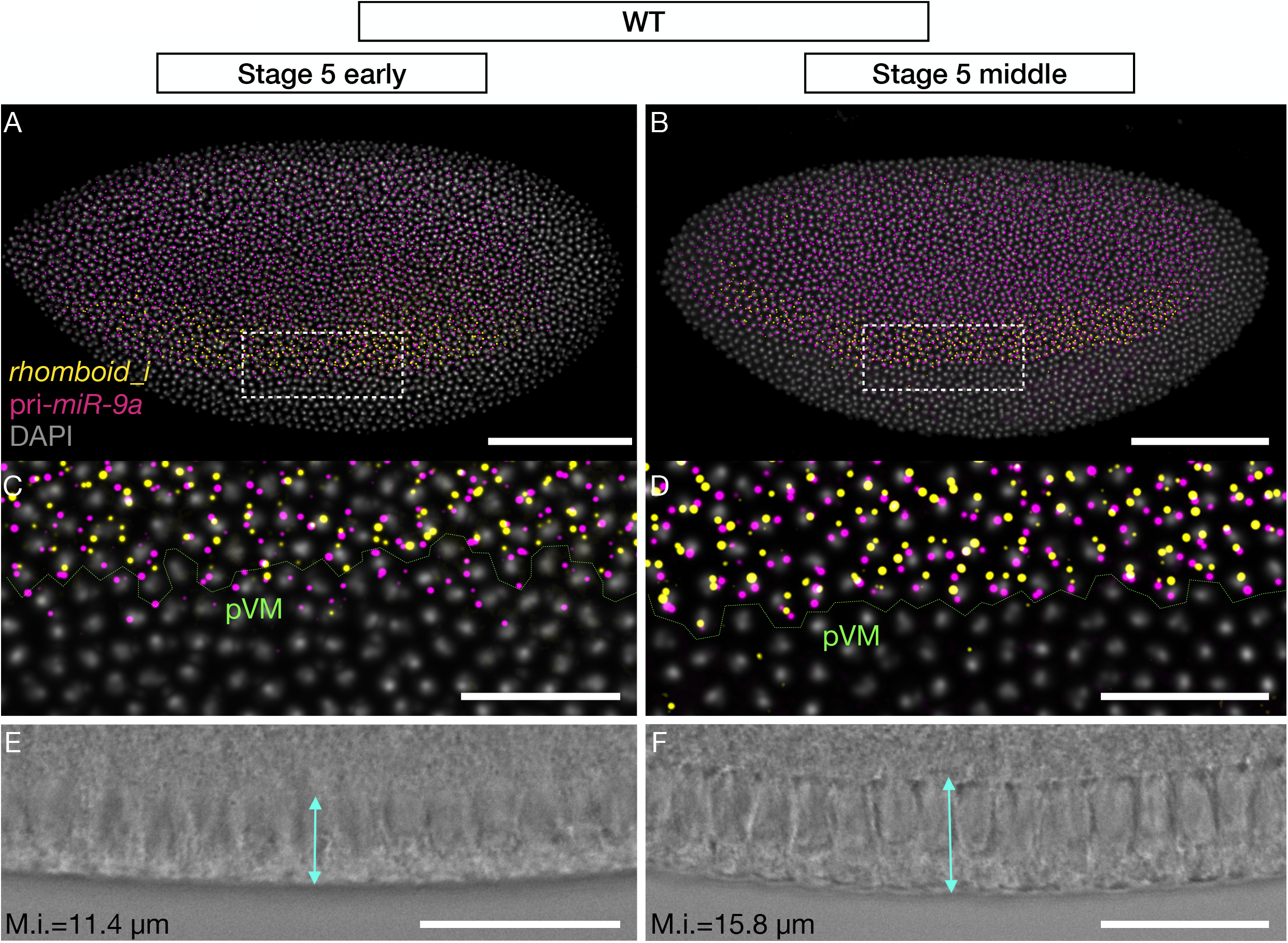
*rhomboid* and *miR-9a* are co-expressed in the neurogenic ectoderm. (A) Early and (B) middle stage 5 *D. melanogaster* embryos stained with probes against *rhomboid* intron (yellow) and the primary transcript of *miR-9a* (magenta). (C-D) zooms of highlighted areas in A and B respectively. In green is highlighted the presumptive ventral midline, which separates mesoderm and ectoderm (pVM). (E-F) Brightfields of ventral borders of the embryos in A and B showing membrane introgression (M.i.). Scalebars: 100 μm (A-B), 25 μm (C-D-E-F).

### Increased *rhomboid* mRNA copy number in *miR-9a*^E39^ mutants

Combining high resolution confocal microscopy with smFISH, immunofluorescence and segmentation allows us to count mRNA molecules in individual cells in *Drosophila* early embryos. We quantified *rho* mRNA molecules per cell in WT and *mir-9a*^E39^ stage 5 embryos (Figure 2 A-B). In order to tightly control the stage of embryonic development, we focused only on stage 5 embryos that have a similar level of membrane introgression. As reported in Fu et al. (2014) the *rho* expression pattern is not spatially or temporally different in *miR-9a* ^E39^ mutant embryos. We imaged and quantified six embryos per genotype and inspected many more and we never saw an abnormal *rho* expression pattern. Nevertheless, when we performed single cell segmentation and quantification, differences started to emerge (see Figure 2 E and F). The data show that the 2 embryos have a spatially equivalent *rho* expression pattern, but the mRNA number per cell is higher in *miR-9a*^E39^ mutant embryos. To corroborate this observation, we performed two independent smFISH experiments using different fluorophores (Figure 2, G-H), with 3 embryos per genotype. The number of cells that have low or no detected *rho* expression varies from embryo to embryo, likely due to stochastic leaky transcription or false positive detection and counting. After excluding cells with fewer than 10 counted *rho* mRNAs, we found that in both experiments, *miR-9a*^E39^ mutants possess a higher number of *rho* mRNA per cell.

**Figure 2.**
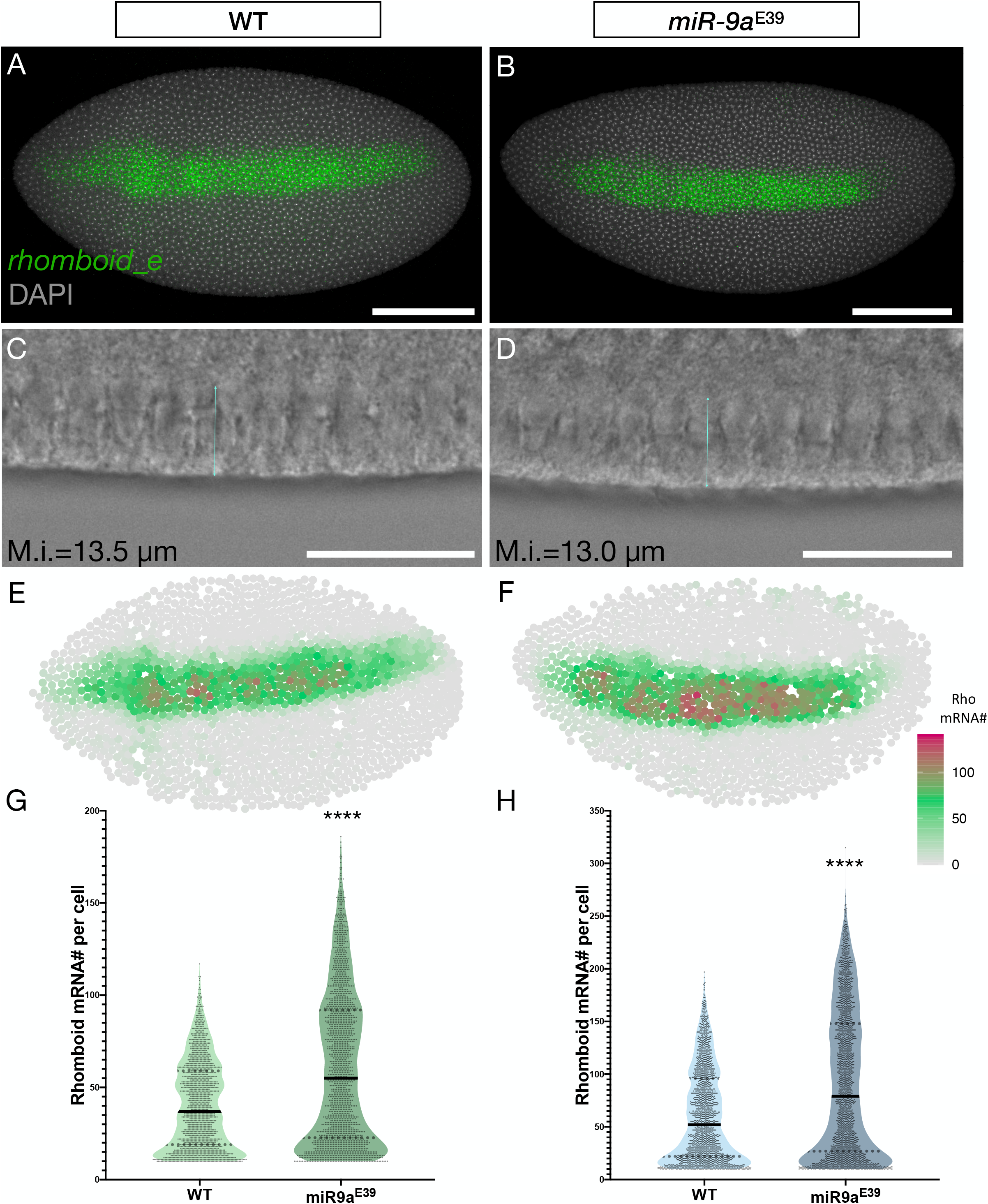
*rhomboid* mRNA number per cell is higher in *miR-9a*^E39^ embryos. (A) WT and (B) *miR-9a*^E39^ middle stage 5 embryos stained with a probe set against *Rhomboid* transcripts. (C-D) Brightfields of a ventral region from embryos in A and B respectively showing membrane introgression. (E-F) Computational reconstruction after segmentation of the embryos in A and B. The colormap is based on mRNA number per cell with grey being low, green intermediate and purple high. (G-H) Two independent quantifications of *rhomboid* mRNA number in single cells in WT and *miR-9a*^E39^ mutant embryos. Each quantification was performed using 3 embryos per genotype. Both p-values <0.0001. Scalebars: 100 μm (A-B), 25 μm (C-D).

To further characterize the difference in *rho* mRNA number in single cells, we coupled the intronic probes used in Figure 1 against *rho* introns with the probes used in Figure 2 against the mature *rho* transcripts, in order to simultaneously quantify *rho* TSs and mature mRNA molecules (Figure 3). We used 100X images to separate and quantify *rho* TS number per cell (maximum 2 per cell prior to replication and 4 per cell following). As the higher magnification does not permit imaging of entire embryos, we focused on the central region of the *rho* expressing stripe, again in stage 5 embryos with a similar membrane introgression (Figure 3 A-B-C, A’-B’-C’). The comparison of *rho* mRNA distribution between WT and *miR-9a*^E39^ embryos again shows that *miR-9a*^E39^ embryos have higher levels of *rho* mRNA number per cell (Figure 3 E). The detection and quantification of *rho* TSs allowed us to distinguish between cells that are differentially transcribing *rho*, and thus subgroup them in 3 classes: cells with no TSs, cells with one TS and cells with two (or more) TSs. In Figure 3-F we reported that cells with a higher number of TSs also show an increased number of *rho* mRNAs, and for each group of cells *miR-9a*^E39^ have a generally higher number of transcripts with respect to WT embryos. This becomes particularly evident for cells that are not transcribing *rho* at the moment the embryo was fixed. It is important to note that very few cells have 3 or 4 TSs (<10 per image over ∼700 segmented cells). These may represent cells following DNA replication, or errors in the segmentation process. We are confident that these small numbers do not significantly affect our analysis.

**Figure 3.**
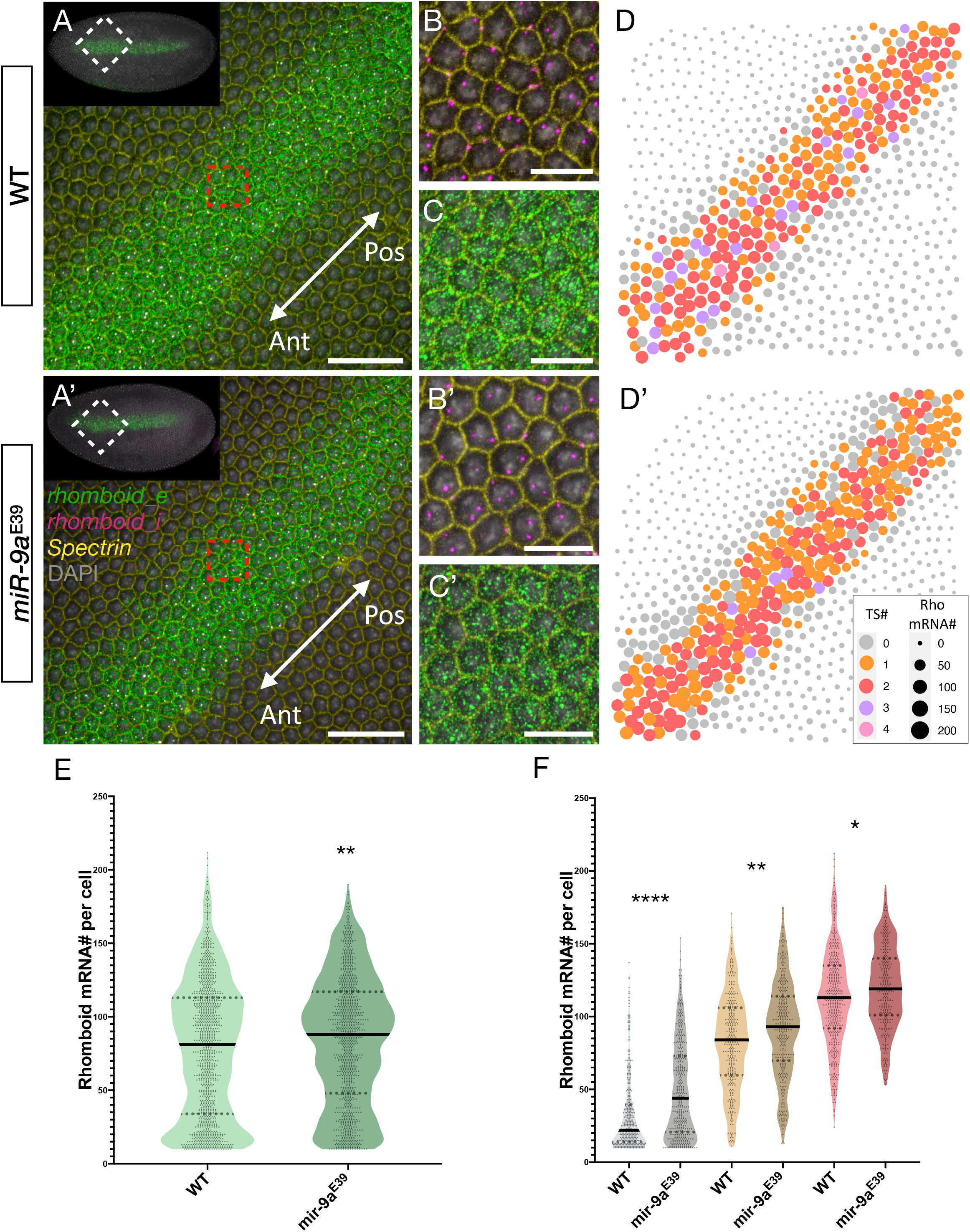
Detection and quantification of *rhomboid* transcription sites in single cells. Central region of (A) WT and (A’) *miR-9a*^E39^ embryos respectively. Orientation is indicated by the white arrow (Ant = Anterior embryonic region, Pos = Posterior embryonic region). (B-B’) Zoom from red area highlighted in A and A’ respectively showing staining against *rhomboid* intron (rhomboid_i, magenta), Spectrin to mark cellular membrane (yellow) and DAPI (grey). (C-C’) Zoom from red area highlighted in A and A’ respectively showing staining against *rhomboid* exon (rhomboid_e, green), Spectrin and DAPI. (D-D’) Computational reconstructions of the images in A and A’ respectively. Each dot corresponds to a segmented cell. The size of the dot corresponds to the number of *rhomboid* mRNAs detected with rhomboid_e, while the colour corresponds to the number of detected transcription sites with rhomboid_i. (E) Comparison between WT and *miR-9a*^E39^ *rhomboid* mRNA number per cell. p-value = 0.0014. (F) Quantified cells are grouped depending on how many alleles are actively transcribing the *rhomboid* locus: grey = 0 alleles active (p-value <0.0001), orange = 1 allele active (p-value = 0.0021), red = 2 or more alleles active (p-value = 0.0259). Scalebars: 100 μm (A-A’), 25 μm (B-C-B’-C’).

### *miR-9a* does not affect cell-to-cell variation in *rhomboid* mRNA number

MicroRNAs are generally thought to have subtle effects on gene expression, mostly acting as buffering factors against intrinsic and extrinsic noise. We therefore investigated whether *miR-9a* might not only affect the number of *rho* transcripts per cell, but also cell-to-cell variability in the number of mature mRNAs present. In order to quantify these effects, we identified the immediate cell neighbours of each segmented cell, and then calculated how variable the *rho* mRNA number per cell is amongst the identified neighbours. As variance scales with mean, areas with high variance do not necessarily correspond to areas in which the cell-to-cell variability is intrinsically higher. Other statistical parameters that have been widely used in order to describe cell-to-cell variability are the coefficient of variation (CV) and the Fano factor (FF) (Munsky *et al*. 2012; Foreman and Wollman 2020). FF is defined as variance/mean while CV as standard deviation/mean. Thus, both measures are mean-normalized. CV is a unitless parameter, and has been used to compare cell-to-cell variability between mRNAs or protein levels resulting from the expression of different genes (Foreman and Wollman 2020). On the other hand, FF has a dimension, and has been used to measure how the observed data are dispersed from a Poisson distribution, which has FF equal to 1 (Thattai and Van Oudenaarden 2001; Hortsch and Kremling 2018). We therefore calculated the FFs for the *rho* mRNA and TS counts reported in Figure 2 and Figure 3 (see Figure 4). We observe that the FF is marginally higher in *miR-9a*^E39^ mutants. Closer inspection shows that the FF is higher in *miR-9a*^E39^ mutants only in the group of cells with no transcription sites, while groups of cells that have a single TS and 2 or more TSs have higher FF in the WT. We speculate that the *miR-9a* buffering action on *rho* mRNA number per cell becomes more evident and/or necessary in quiescent cells that are not actively transcribing *rho*.

**Figure 4.**
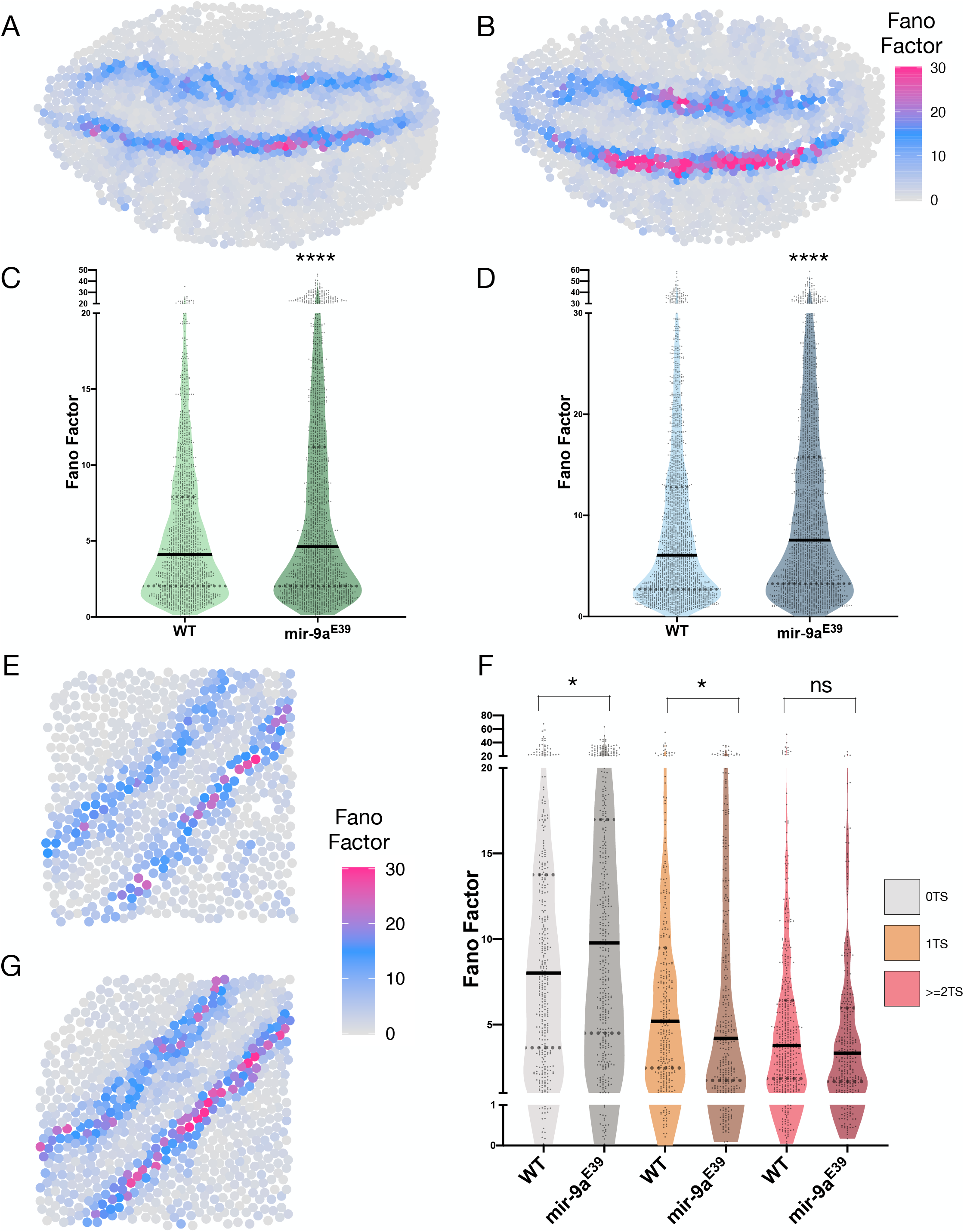
Fano factor quantification and comparison between WT and *miR-9a*^E39^ mutant embryos. Computational reconstruction of Fano factor distribution calculated in neighbour clusters in (A) WT and (B) *miR-9a*^E39^ stage 5 embryos. These two embryos are the same reported in figure 2 E-F respectively. (C-D) Comparison between Fano factor in WT and *miR-9a*^E39^ embryos in 2 indepndent experiments (n = 3 embryos each). P-value < 0.0001 in both graphs. (E-G) Graphical reconstruction of Fano factor distribution calculated in neighbour cells clusters in a WT and *miR-9a*^E39^ embryos, corresponding to Figure 3 A-A’ respectively. (F) cells are sub-grouped depending on their transcription sites number. p-values = 0.0147 (0 TS) and 0.0123 (1 TS), ns = non-significant.

### Rho is over-expressed in *miR-9a*^E39^ mutants during embryonic stage 5 and 6

As a change in mRNA levels does not necessarily linearly corelate with the change in accumulation of the encoded protein (Koussounadis *et al*. 2015), we compared Rho protein levels between WT and *miR-9a*^E39^ embryos. It has been reported that Rho protein expression is detectable from the embryonic stages 10-11 in WT animals, despite *rho* mRNA being transcribed much earlier during stage 5 (Llimargas and Casanova 1999). However, we find that during stage 5, Rho protein was detectable in *miR-9a*^E39^ embryos. In Figure 5 we show Rho staining in stage 5 and stage 6 WT and *miR-9a*^E39^ embryos with relative quantifications. Anti-Dorsal antibody was used to provide a further control on the quality of the staining and to orient the embryos. Fluorescence measurements were performed in FIJI by randomly selecting 15 areas per embryo (5 in the anterior, 5 in the central and 5 in the posterior regions). Quantifications shown in Figure 5 (panels C and F for stage 5 and 6 respectively) clearly show that Rho levels are significantly higher (p-value < 0.0001 in both cases) in *miR-9a*^E39^ mutants.

**Figure 5.**
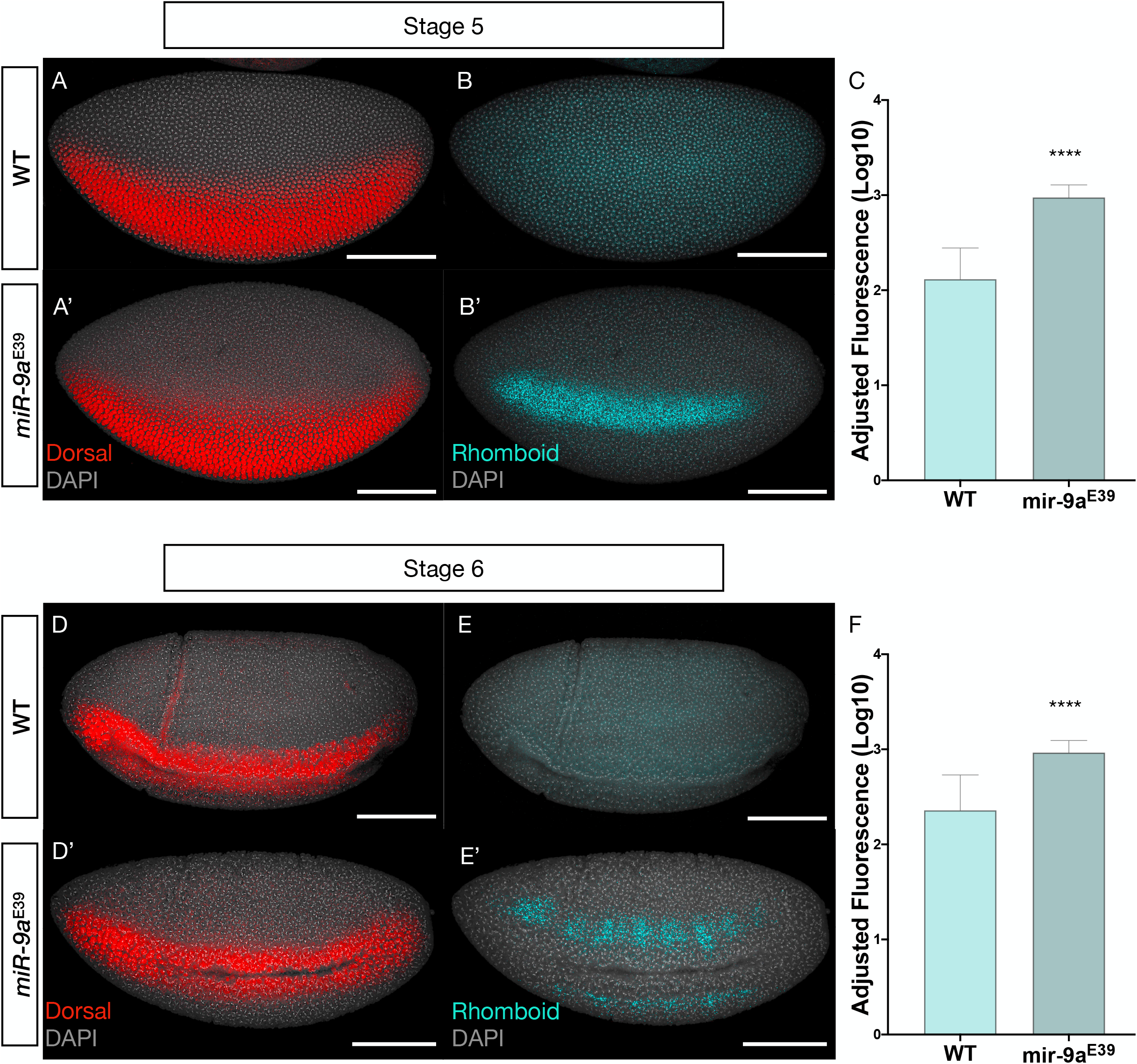
Rhomboid protein is over-expressed in *miR-9a*^E39^ embryos during stage 5 and 6. (A-B) Stage 5 WT and (A’-B’) *miR-9a*^E39^ embryos respectively stained against Dorsal (red) and Rhomboid (cyan). (C) Adjusted fluorescence levels from Rhomboid staining in stage 5 embryos (n=3 per genotype). In each embryo 15 areas equally distributed along the Dorsal expression border were quantified. Measurements are reported in Log10 scale. P-value < 0.0001. (D-E, D’-E’) Stage 6 WT and *miR-9a*^E39^ embryos respectively stained against Dorsal (red) and Rhomboid (cyan). (F) Adjusted fluorescence levels from Rhomboid staining in stage 6 embryos (n=3 per genotype). Quantified as in (C). P-value < 0.0001. Scalebars: 100 μm in all panels.

## Discussion

*rho* has been one of the most studied Dorsal target genes. Its expression becomes restricted to the neurogenic ectoderm in a precisely orchestrated manner: the low nuclear levels of Dorsal in the dorsal ectoderm do not support *rho* activation, while *snail* represses its transcription in the mesoderm (Ip *et al*. 1992b; Hong *et al*. 2008). *rho* has not been previously studied as a direct target of miRNA regulation, but the combined effect of mutations in *miR-1* and *miR-9a* on *rho* mRNA distribution motivated our investigation into *rho* regulation by miRNAs (Fu *et al*. 2014). We found that the per cell copy number of *rho* mRNA is significantly higher in *miR-9a* ^E39^ mutant embryos (Figure 2 and Figure 3), suggesting *miR-9a* affects *rho* mRNA stability or degradation. We could not find a clear effect of *miR-9a* on cell-to-cell variability of the number of either *rho* mRNA transcription sites or mRNA molecules (Figure 4). Nevertheless, when we distinguish between cells that are and are not actively transcribing *rho*, we find that the FF of cells with no transcription sites was significantly higher in *miR-9a*^E39^ mutants. This leads us to suggest that, in WT animals, *rho* mRNA is “rapidly” degraded when transcription stops, whereas this degradation is less efficient when *miR-9a* is removed, and cell heterogeneity consequently increases. To our knowledge, this is the first study in which mRNA copy number was compared in different genotypes using single cell quantitative microscopy in order to uncover miRNA regulatory roles on target gene expression.

It has been shown that protein levels are usually more stable than mRNA levels (Perl *et al*. 2017). The *miR-9a* regulatory effect on Rho protein accumulation might therefore be more evident than the one we observed on the mRNA as it better reflects the integrated activity over time. Rho is a transmembrane protease localized in the Golgi. While Fu et al. reported *rho* mRNA patterns in double *miR-9a*/*miR-1* mutants (Fu *et al*. 2014), no information on the protein pattern was previously available. We observed dramatic differences in timing and level of Rho protein accumulation when comparing WT and *miR-9a*^E39^ embryos. In the WT, Rho was only detectable from stage ∼10, whereas in *miR-9a*^E39^ embryos it was clearly present from stage 5, the same stage when we see *rho* transcription initiate. The early accumulation of Rho protein appears to be inhibited by *miR-9a*. We suggest that translational inhibition by *miR-9a* is released when a certain level of *rho* mRNA is reached, or in response to an external signal later in development. We also note the possibility that early low levels of Rho protein accumulation may be present but are undetectable with current technology.

Previous work on the *miR-9a*/*miR-1* double mutant shows that when *miR-1* is also removed, greater developmental defects emerge leading to failure of gastrulation and ventral midline enclosure (Fu *et al*. 2014). This phenotype suggests that these two miRNAs play an important role in germ layer differentiation. Indeed, while *miR-9a* and *miR-1* involvement in dorso-ventral (DV) axis patterning has not been definitively established, their expression patterns indicate they are targets of DV specification (Biemar *et al*. 2006). Our current findings provide convincing evidence for a role of *miR-9a* in the DV patterning process during early *Drosophila* embryogenesis. We posit that *miR-9a* regulates *rho* mRNA accumulation and translation, possibly affecting Epidermal Growth Factor Receptor (EGFR) signalling and specification of the dorsal and neurogenic ectoderm (Golembo *et al*. 1996; Guichard *et al*. 1999). The role of *miR-1* is less clear as *miR-1* is not expressed in the same region as *rho*, and therefore *miR-1* can affect *rho* expression only indirectly. *miR-1* is involved in muscle development and is exclusively expressed in the mesoderm (Sokol and Ambros 2005). We suggest that the combination of disrupted *miR-1* function in the mesoderm and *miR-9a* function in the neurogenic ectoderm leads to disruption in establishment or maintenance of an organized border between these two germ layers, as seen in the double mutants (Fu *et al*. 2014).

To conclude, we have shown in this work a new role of a the well conserved *miR-9a* during early *Drosophila* embryogenesis. We have observed that *miR-9a* affects both *rho* mRNA copy number per cell (possibly by degradation) and inhibits *rho* translation. Our findings also show the importance of single-cell quantification when studying the effects of miRNA regulation on target genes. As miRNAs act as weak modulators of gene expression, single-cell quantitative approaches can reveal previously unknown effects on mRNA and protein regulation by miRNAs. This work and the methods described can be easily applied to many other miRNA-target gene networks to allow new insights into miRNA function during development.

## Acknowledgments

We thank the staff from the University of Manchester’ Bioimaging Facility, in particular Dr. Peter March, for help with confocal microscopy. We also thank the Hayashi lab for providing anti-Rho antibodies, the Fen-Biao Gao lab for providing *miR-9a*^E39^ flies and Dr. Fabian Morales-Polanco for discussions and comments on the manuscript.

## Author contributions

LG, MR and SGJ conceived the project. Experiments were designed by LG and MR and performed by LG. The manuscript was written by all the authors.

## Supplementary material

**Table S1.**
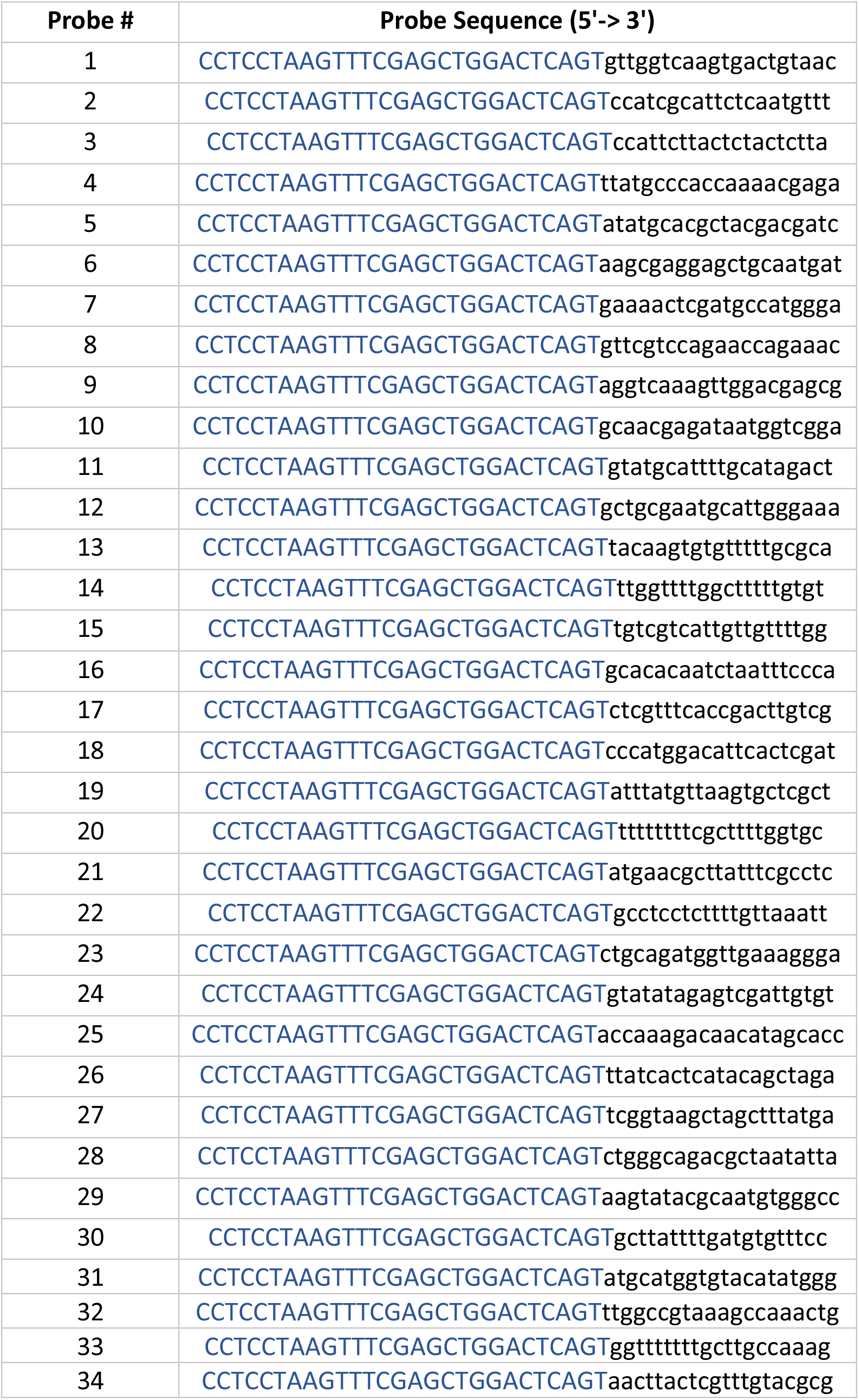
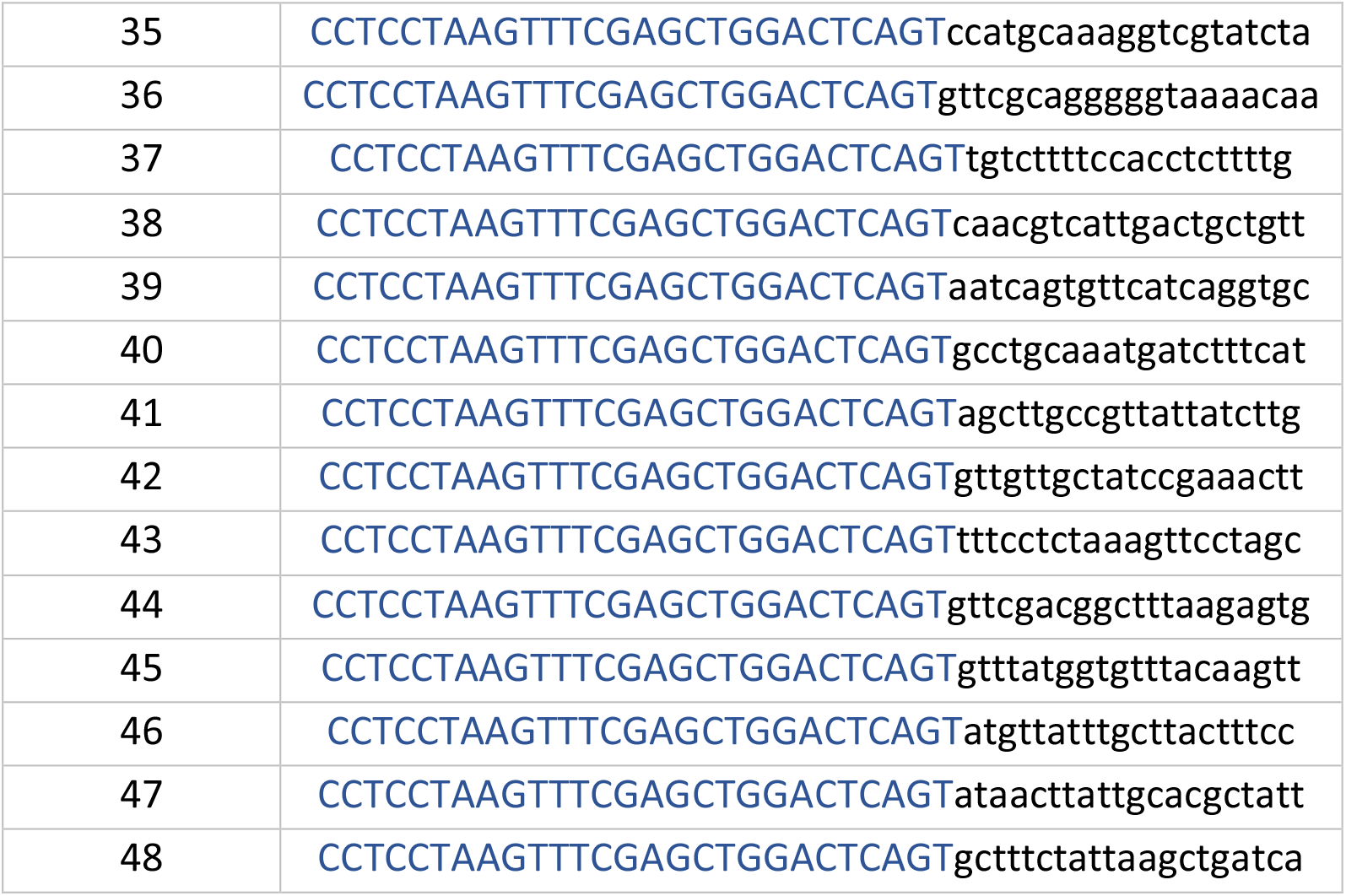
Probes against *mir-9a*

**Table S2.**
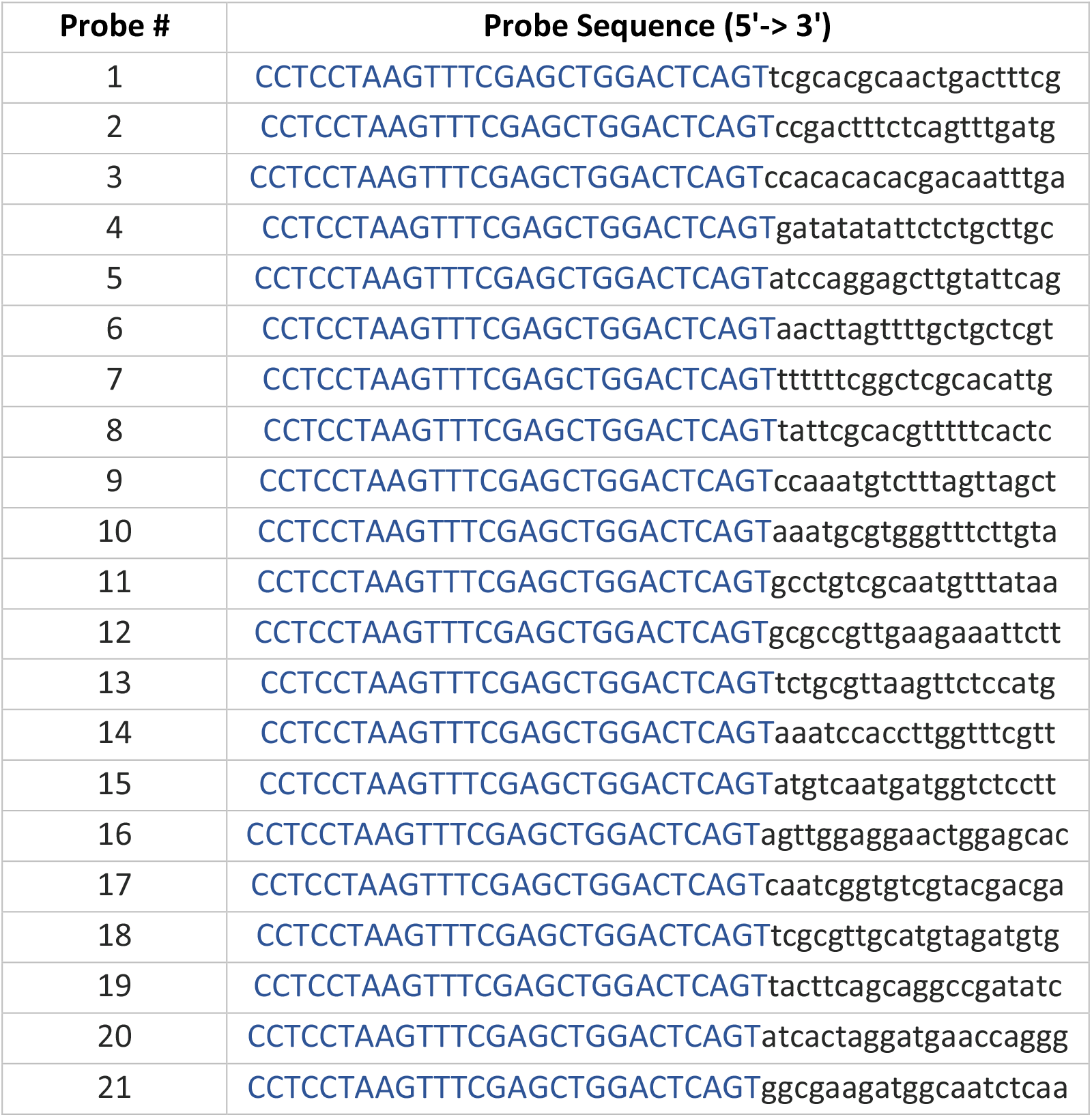
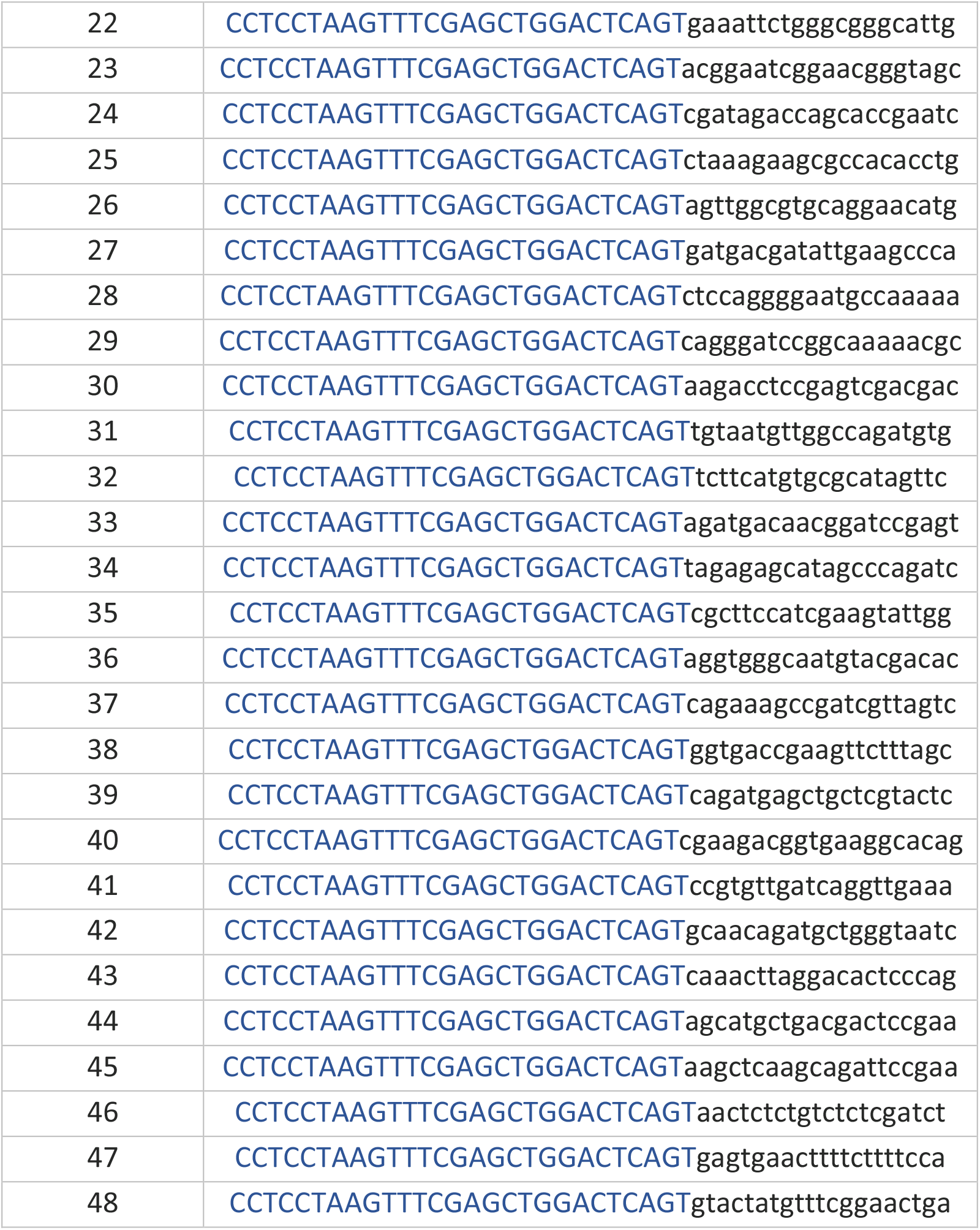
Probes against r*homboid* exons (rhomboid_e)

**Table S3.**
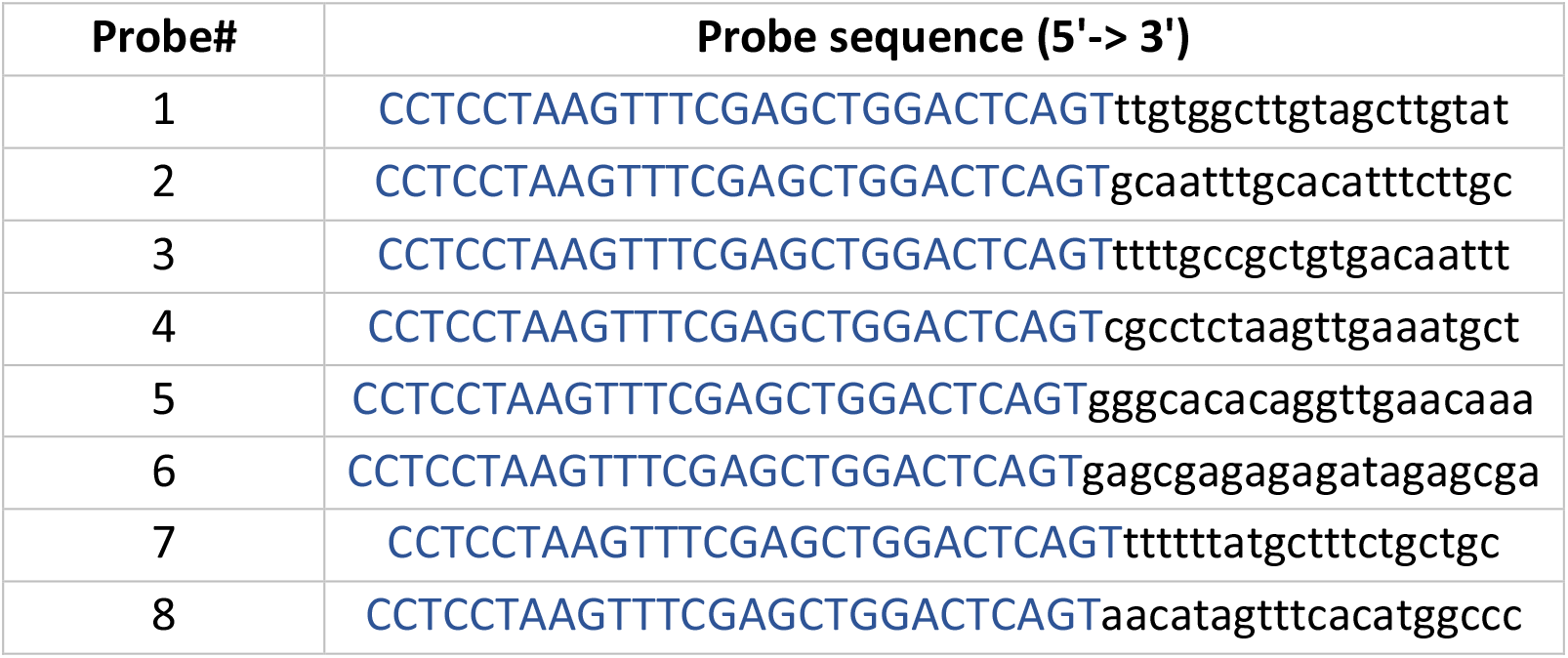
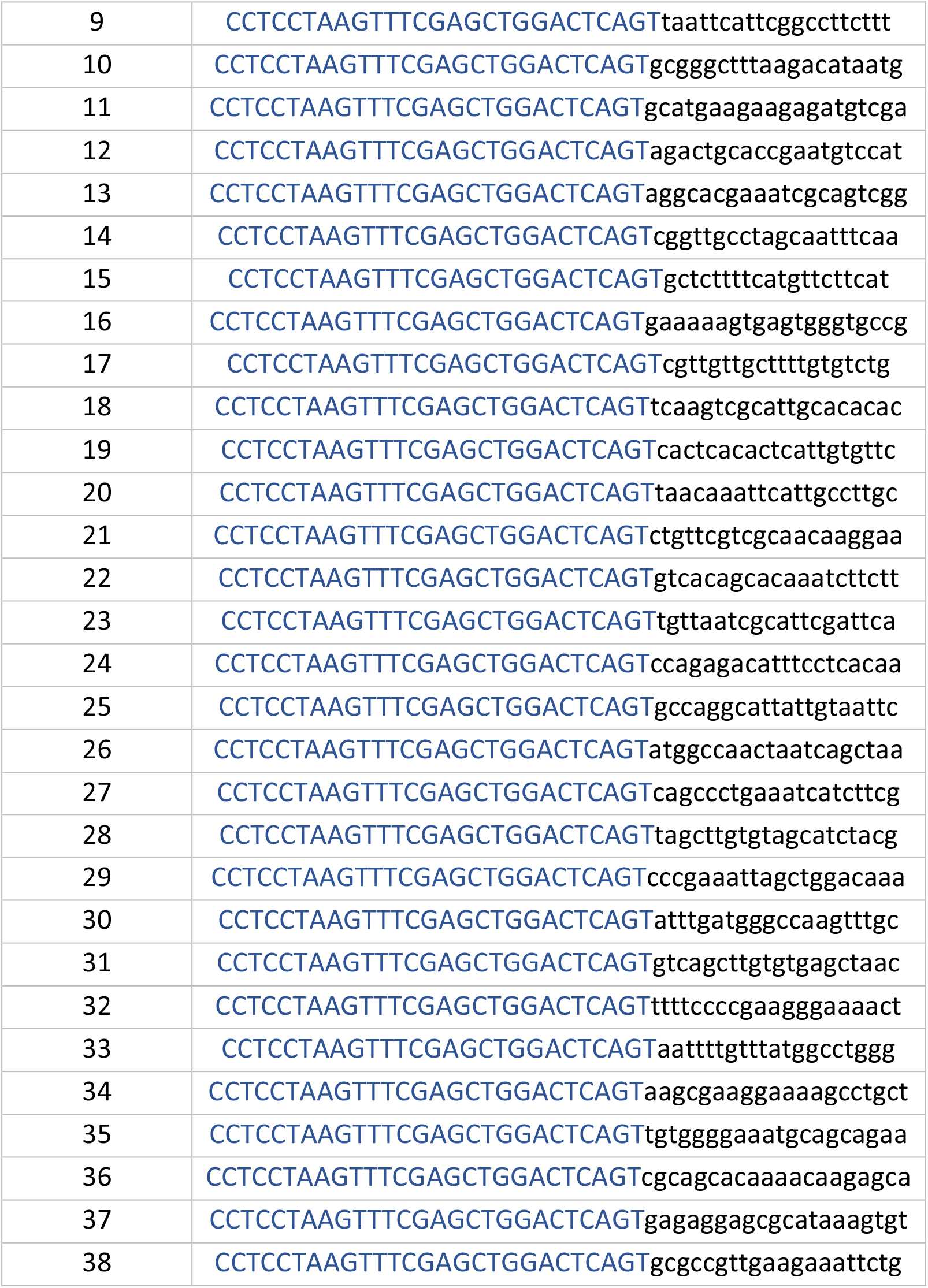
Probes against *rhomboid* intron (rhomboid_i)

